# Joint annotation of chromatin state and chromatin conformation reveals relationships among domain types and identifies domains of cell type-specific expression

**DOI:** 10.1101/009209

**Authors:** Maxwell W. Libbrecht, Ferhat Ay, Michael M. Hoffman, David M. Gilbert, Jeffrey A. Bilmes, William Stafford Noble

## Abstract

The genomic neighborhood of a gene influences its activity, a behavior that is attributable in part to domain-scale regulation, in which regions of hundreds or thousands of kilobases known as domains are regulated as a unit. Previous studies using genomics assays such as chromatin immunoprecipitation (ChIP)-seq and chromatin conformation capture (3C)-based assays have identified many types of regulatory domains. However, due to the difficulty of integrating genomics data sets, the relationships among these domain types are poorly understood. Semi-automated genome annotation (SAGA) algorithms facilitate human interpretation of heterogeneous collections of genomics data by simultaneously partitioning the human genome and assigning labels to the resulting genomic segments. However, existing SAGA methods can incorporate only data sets that can be expressed as a one-dimensional vector over the genome and therefore cannot integrate inherently pairwise chromatin conformation data. We developed a new computational method, called graph-based regularization (GBR), for expressing a *pairwise prior* that encourages certain pairs of genomic loci to receive the same label in a genome annotation. We used GBR to exploit chromatin conformation information during genome annotation by encouraging positions that are close in 3D to occupy the same type of domain. Using this approach, we produced a comprehensive model of chromatin domains in eight human cell types, thereby revealing the relationships among known domain types. Through this model, we identified clusters of tightly-regulated genes expressed in only a small number of cell types, which we term “specific expression domains.” We additionally found that a subset of domain boundaries marked by promoters and CTCF motifs are consistent between cell types even when domain activity changes. Finally, we showed that GBR can be used for the seemingly unrelated task of transferring information from well-studied cell types to less well characterized cell types during genome annotation, making it possible to produce high-quality annotations of the hundreds of cell types with limited available data.

## Introduction

Although the mechanism of regulation of a gene by a promoter directly upstream of its transcription start site is well understood, this type of local regulation does not explain the large effect of genomic neighborhood on gene regulation. The neighborhood effect is in part the consequence of domain-scale regulation, in which regions of hundreds or thousands of kilobases known as domains are regulated as a unit (Bickmore and van Steensel, 2013; Chakalova et al., 2005; Akhtar et al., 2013). Current understanding of domain-scale regulation is based on a number of domain types, each defined based on a different type of data, such as histone modification ChIP-seq, replication timing, or measures of chromatin conformation. However, as a result of the difficulty of integrating genomics data sets, the relationships among these domain types are poorly understood. Therefore, a principled method for jointly modeling all available types of data is needed to improve our understanding of domain-scale regulation.

A class of methods we term *semi-automated genome annotation* (SAGA) algorithms are widely used to jointly model diverse genomics data sets. These algorithms take as input a collection of genomics data sets and simultaneously partition the genome and label each segment with an integer such that positions with the same label have similar patterns of activity. These algorithms are “semi-automated” because a human performs a functional interpretation of the labels after the annotation process. Examples of SAGA algorithms include HMMSeg (Day et al., 2007), ChromHMM (Ernst and Kellis, 2010), Segway (Hoffman et al., 2012) and others (Thurman et al., 2007; Lian et al., 2008; Filion et al., 2010). These genome annotation algorithms have had great success in interpreting genomics data and have been shown to recapitulate known functional elements including genes, promoters and enhancers.

However, existing SAGA methods cannot model chromatin conformation information. The 3D arrangement of chromatin in the nucleus plays a central role in gene regulation, chromatin state and replication timing (Misteli, 2007; Dekker, 2008; Ryba et al., 2010; Dixon et al., 2012). Chromatin architecture can be investigated using chromatin conformation capture (3C) assays, including the genome-wide conformation capture assay, Hi-C. A Hi-C experiment outputs a matrix of contact counts, where the contact frequency of a pair of positions is inversely proportional to the positions’ 3D distance in the nucleus (Lieberman-Aiden et al., 2009; Ay et al., 2014b). Existing SAGA methods can incorporate any data set that can be represented as a vector defined linearly across the genome, but they cannot incorporate inherently pairwise Hi-C data without resorting to simplifying transformations such as principle component analysis.

We present a method for integrating chromatin architecture information into a genome annotation method. Motivated by the observation that pairs of loci close in 3D tend to occupy the same type of domain, we encourage these pairs to be assigned the same label in a genome annotation through a *pairwise prior.* We developed a novel computational method, called graph-based regularization (GBR), which performs inference in the presence of such a pairwise prior, and we extended the existing SAGA algorithm Segway (Hoffman et al., 2012) to implement this method.

We found that integrating Hi-C data into Segway using GBR resulted in annotations which predict replication time and self-interacting domains known as topological domains (Dixon et al., 2012) better than annotation without GBR. We then applied Segway with GBR to produce a comprehensive model of domain-scale regulation in the human lung muscle fibroblast cell line IMR90. This model revealed a set of five domain types which capture the known properties of existing domain types and thereby reveal the relationships among these types. Using this model, we identified domains characterized by high regulatory activity and genes specifically expressed in a small number of cell types, which we term “specific expression domains.” We additionally annotated seven other cell types and discovered that domain boundaries are often consistent between cell types even when domain state changes, a trend which can be explained in part by the presence of promoters and CTCF motifs.

GBR can also be used for the seemingly-unrelated task of transferring information from well-studied cell types for the annotation of cell types with limited available data. Consortia such as ENCODE have characterized a small set of cell types in great detail using hundreds of genomics assays. However, due to the high cost of genomics experiments, it is feasible to perform only a few assays on any additional cell type of interest. For example, ENCODE and Roadmap Epigenomics have each performed 2-10 experiments in more than 100 cell types, and it is common for an individual lab to perform a small number of experiments on a particular cell type or perturbation of interest. In such settings, it is crucial to leverage information garnered from well-studied cell types to allow accurate annotation of other cell types using just a few experiments. We transfer information from well-studied cell types with GBR by using the pairwise prior that loci that were assigned the same label in many well-studied cell types should be more likely to receive the same label in a cell type of interest. We demonstrate the efficacy of this method by holding out data in the annotation of a well-studied cell type and showing that using GBR to incorporate information from a related cell type improves the accuracy with which the annotation predicts functional elements over annotation without GBR. Therefore, GBR makes it possible to produce high-quality annotations of the hundreds of cell types with limited available data.

## Results

### Chromatin domains co-localize with domains of similar activity

Previous research has shown that large chromatin domains (~1 MB) tend to co-localize with domains of similar activity in 3D (Ryba et al., 2010; Lieberman-Aiden et al., 2009). To further explore this trend, we compared the 1D genomics data at pairs of statistically significantly interacting loci (Supplementary Figure 1). As expected, we found that chromatin signals were highly consistent at pairs of positions nearby in 3D. First, histone modification and replication signal values at pairs of significantly interacting loci are more highly correlated than for a rotational permutation control, which controls for the 1D pattern of the signal (Supplementary Figure 1A). Second, Segway labels generated without using GBR from an annotation of the genome were assigned the same label much more often than a rotational permutation control (Supplementary Figure 1B). Note that this pattern is in stark contrast to small elements (~ 100bp) such as promoters and enhancers, which do not, in general, cluster with elements of the same type. These observations suggest that chromatin conformation data might best be incorporated using a pairwise prior stating that a pair of positions should be more likely to receive the same label if the positions are close in 3D. Therefore, we sought to develop new methods for leveraging chromatin conformation data using this co-localization pattern.

### Graph-based regularization expresses a pairwise prior in a SAGA method

Existing SAGA algorithms use dynamic programming algorithms to perform inference in a chain-structured Bayesian network such as a hidden Markov model. Dynamic programming algorithms such as the forward-backward algorithm can be used to perform inference efficiently in models with chain-structured dependencies; however, applying these methods in the presence of a pairwise prior that connects arbitrary pairs of positions results in inference costs which grow exponentially in the number of genomic positions. Therefore, these methods cannot be applied to genome annotation problems with millions or billions of variables. We propose a novel convex optimization framework which allows for efficient inference in this case.

The method takes as input a set of genomics data sets and weighted graph over the genomic positions, where a large weight on a given pair of positions indicates that we have a strong prior belief that this pair should receive the same label. It outputs a probability distribution over the integer labels at each position. The method encourages pairs of positions connected by edges in the graph to be assigned the same label by minimizing a measure of dissimilarity between their output probability distributions called the Kullback-Leibler (KL) divergence.

We call this strategy of using a graph to incorporating a pairwise prior *graph-based regularization* (GBR) (Methods). Note that in this manuscript we use the word “prior” in the non-technical sense of “prior information,” not in the sense of a prior distribution for a Bayesian model. We have developed an efficient, novel alternating minimization algorithm that optimizes this objective (Supplementary Note 1). Using synthetic data, we determined that graph-based regularization outperforms alternative methods based on approximate inference as well as existing methods for graph-based regularization (Supplementary Note 2). We then extended the SAGA method Segway to implement this algorithm (Hoffman et al., 2012) (Supplementary Note 3).

### Using GBR to integrate 3D structure information improves prediction of replication and topological domains

We used graph-based regularization to integrate chromatin conformation information using the pairwise prior that positions close in 3D should be more likely to be identified as the same domain type (Figure 1A). To do this, we construct a GBR graph that connects each pair of positions with weight proportional to our statistical confidence that the positions physically interact (Ay et al., 2014a) (Methods). This measure of statistical confidence controls for the bias of Hi-C for positions close in 1D as well as biases for sequence features such as GC content and restriction site density.

**Figure 1:**
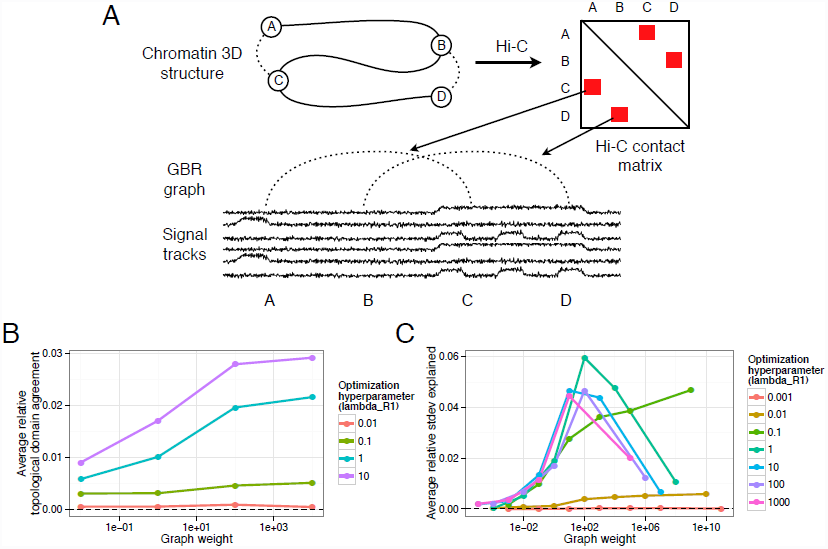
(A) Strategy for incorporating Hi-C data using GBR. (B) Effect of GBR hyperparameters on topological domain agreement. The X axis indicates the value of the graph weight hyperparameter *λ*_*G*_. Y axis indicates the average over 10 annotations (one for each input histone modification data set) of the fraction improvement in topological domain label agreement by adding GBR over Segway without GBR. (C) Same as (B), but Y axis indicates improvement in replication timing standard deviation explained and average is over 29 annotations. Annotations used four labels, 10kb resolution.

Incorporating Hi-C data using GBR has the effect of both aligning domains to regions of self-interacting chromatin and helping to determine the label of each segment. We evaluated the first effect, as a sanity check, by computing the accuracy with which our annotation predicts self-interacting regions of chromatin of size ~ 1 MB called topological domains (Dixon et al., 2012). We evaluated the second effect by comparing to replication time, which is highly correlated with gene expression and chromatin state and therefore is a good proxy for domain type. In order to evaluate our performance in a variety of conditions, we ran Segway augmented with GBR separately once for each of the 29 histone modification sets available in IMR90, in each case using as input a single histone modification data set and a GBR graph based on IMR90 Hi-C data. We found that the annotation’s ability to identify both topological domains (*p <* 10^-16^, t-test) and replication time (*p* < 10^-16^, t-test) was greatly improved by using GBR (Figure 1B,C). To evaluate the degree to which an annotation matches topological domains, we computed, for each topological domain, the fraction of positions in the topological domain receiving the same label, controlling for the length and label distribution by comparing to a circularly permuted annotation (Methods). We measure the degree to which an annotation predicts replication time by the variance in replication timing explained by the annotation (Methods). The improvement from adding Hi-C with GBR was greater than the improvement achieved by instead adding another histone modification data set, for 26/30 and 27/30 histone modification data sets for topological domains and replication time, respectively. Furthermore, this improvement was consistent for a large range of hyperparameters (several order of magnitude around optimal, Figure 1B,C). These results demonstrate that incorporating Hi-C data using GBR greatly improves the quality of the resulting annotation, and moreover that Hi-C is more informative for determining domain identity than most other data types.

### Joint domain annotation of chromatin state and chromatin conformation captures previously-described domain types

Having verified the utility of GBR for incorporating Hi-C data, we next sought to investigate domain-scale genome regulation using this method. Current understanding of domain-scale regulation is based on a number of domain types (seven, by our count), each defined based on a different type of data, such as histone modification, replication timing or 3C-based assays (Table 1). For example, ChIP-seq on the histone modification H3K27me3 has revealed repressive domains known as facultative heterochromatin (Morey and Helin, 2010; Pauler et al., 2009), and 3C-based assays have revealed regions of self-interacting chromatin known as topological domains (Dixon et al., 2012). Because all of these domain types are defined using different types of data, until now it has been difficult to understand the relationships among these domain types. GBR provides a principled method for integrating all types of data into a unified annotation of domains. We therefore used Segway with GBR to create such a unified annotation, in order to understand what types of domains exist and their interrelationships.

**Table 1:** Known types of domains. Note that the lengths of domains depend greatly on the method used to define them.

| Name | Label(s) | Typical length | Relevant data types | References |
| --- | --- | --- | --- | --- |
| Topological | All | 0.8 MB | Hi-C | Dixon et al. (2012) |
| Open compartment / early replicating | BRD, SPC, FAC | 10 MB | Hi-C, Repli-(chip/seq) | Lieberman-Aiden et al. (2009)<br>Ryba et al. (2010) |
| Closed compartment / late replicating | QUI, CON, FAC | 10 MB | Hi-C, Repli-(chip/seq) | Lieberman-Aiden et al. (2009)<br>Ryba et al. (2010) |
| Quiescent | QUI | 0.5 MB | None | Ernst and Kellis (2010)<br>Hoffman et al. (2012) |
| Constitutive heterochromatin | CON | 0.5 MB | H3K9me3 (ChIP-seq) | Lachner et al. (2003) |
| Facultative heterochromatin | FAC | 0.5 MB | H3K27me3 (ChIP-seq) | Morey and Helin (2010)<br>Pauler et al. (2009) |
| Lamina | See text | 0.6 MB | lamin B1 (DamID) | Guelen et al. (2008)<br>Wen et al. (2009) |
| Broad activity | BRD | 0.5 MB | Transcription (i.e. H3K36me3) (ChIP-seq) | Novel in human cells (see text) |
| Specific activity | SPC | 0.5 MB | Regulation (i.e. H3K27ac) (ChIP-seq) | Novel in human cells (see text) |

We annotated the cell type IMR90 using all 30 signal data sets we had available in IMR90 and a GBR graph derived from IMR90 Hi-C data, resulting in an annotation with median segment length of 0.4 MB (Figure 2A,B,C, Methods, Supplementary Tables 1,2). Including Hi-C into this annotation using GBR changed the label of 6% of positions relative to an annotation without GBR, meaning Hi-C has a slightly larger influence than the 1/31 = 3% difference one would expect from adding one additional data set (Supplementary Figure 5). Because determining the optimal number of labels for an annotation remains an open problem, we specified a somewhat larger number of labels than we expected to be supported in the data (eight), then manually merged labels that we deemed to be redundant.

**Figure 2:**
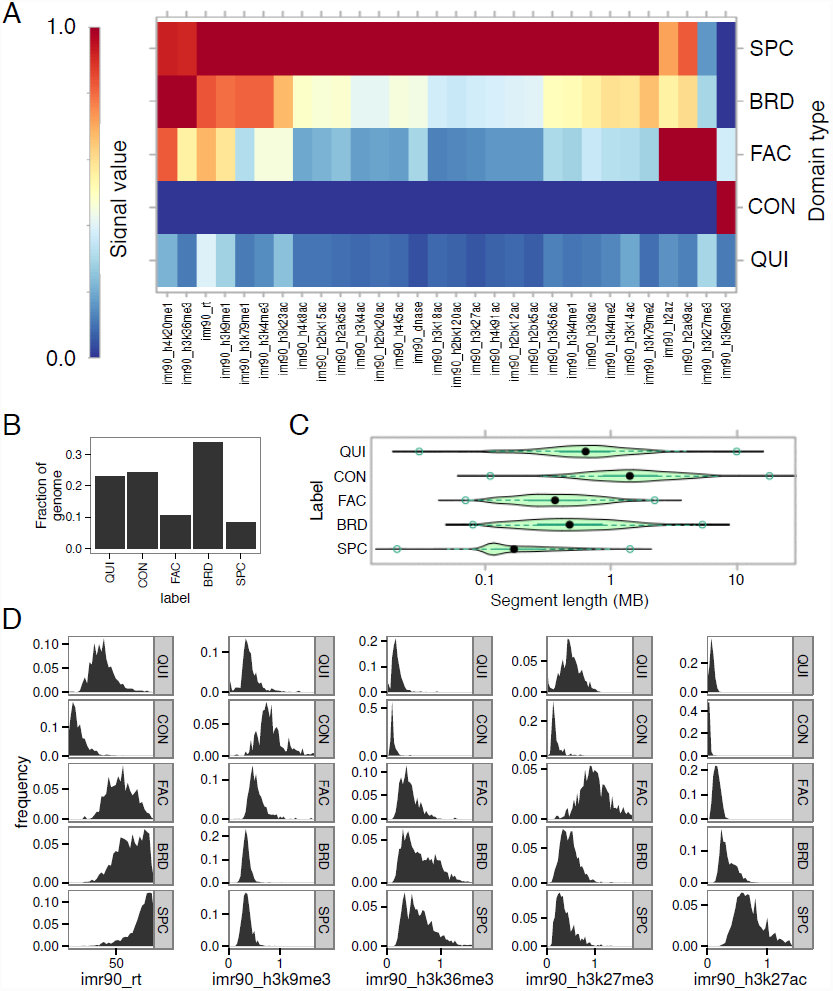
Statistics of domain annotation made with Segway augmented with GBR. (A) Heatmap of association of IMR90 domain labels to IMR90 data sets used in training. (B) Fraction of genome covered by each domain type. (C) Segment length distribution for each domain type. (D) Label-specific histograms of values for five notable data sets in IMR90.

We compared our annotation to eight types of features (Table 2). On the basis of these analyses, we merged labels that appeared redundant and assigned names to each integer label (or group of labels) that best match our interpretation of their function. This procedure yielded five types of domains: (1) broad expression (BRD), (2) specific expression (SPC), (3) facultative heterochromatin (FAC), (4) constitutive heterochromatin (CON), and (5) quiescent (QUI). We describe the analyses that led us to these names in the following sections.

**Table 2:** Summary of learned domain types.

| Data type | Figure | QUI | CON | FAC | BRD | SPC |
| --- | --- | --- | --- | --- | --- | --- |
| Median segment length | 2C | 0.6 MB | 1.4 MB | 0.4 MB | 0.6 MB | 0.2 MB |
| Histone modifications | 2A,D | None | H3K9me3 | H3K27me3 | Transcription (i.e. H3K36me3) | Regulation (i.e. H3K27ac) |
| Replication timing | 2A,D | Switching late | Constitutively late | Mixed | Constitutively early | Switching early |
| GC content | 3D | Low | Low | High | Mid | High |
| Conservation | 3E | Conserved | Nonconserved | Conserved or accelerated | Conserved | Conserved |
| Hi-C eigenvalue compartment | 3F | Closed | Closed | Mixed | Open | Open |
| Gene density | 3A | Low | Low | High | High | High |
| Gene expression | 3B | N/A | N/A | Repressed | Average | Increased |
| Lamin | 4 | Core | Core | Boundaries | Flanks | Flanks |

In order to understand how domains change state between cell types, we additionally annotated eight cell types using twelve data sets present in all eight types and a GBR graph representing common 3D contacts generated by combining IMR90 and H1-hESC Hi-C data sets (Methods, Supplementary Table 2). This strategy of combining Hi-C data sets is motivated by the consistency of Hi-C across cell types (Supplementary Figure 3). Again, we used eight labels and merged redundant labels, to which we assigned the same five names. We investigated the properties of these five domain types.

### Repressive domains are divided into constitutive, facultative and quiescent heterochromatin

Previous studies have reported two types of repressive domains. The first type, best known as “constitutive heterochromatin” but sometimes referred to simply as “heterochromatin”, is regulated by the HP1 complex and associated with the histone modification H3K9me3 (Lachner et al., 2003). Constitutive heterochromatin is thought to repress permanently silent regions such as centromeres and telomeres. As expected, one output domain type “CON” exhibits all the known properties of constitutive heterochromatin. CON domains are associated with H3K9me3 (Figure 2A,D), are extremely depleted for genes (Figure 3A), are associated with low GC content and lack of evolutionary conservation (Figure 3D,E), appear within the Hi-C eigenvector closed compartment (Figure 3F, Methods), and cover regions which are constitutively late replicating in all cell types (Figure 3G). CON domains are depleted both for transcription factor motifs and for transcription factor binding at motifs (Figure 3H, I).

**Figure 3:**
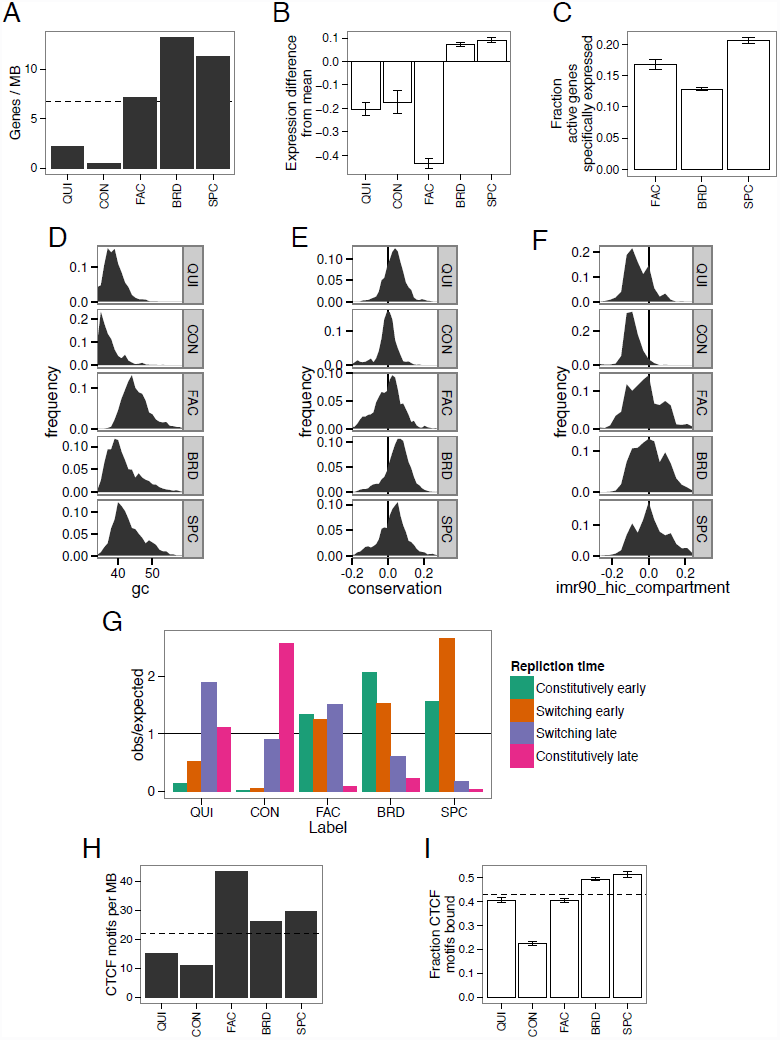
Characteristics of domain types. (A) Gene density in IMR90. (B) Gene expression relative to average over 33 cell types, averaged over eight annotations (t-test 95% confidence interval error bars). (C) Fraction of active genes also active in more than 15 other cell types, averaged over eight annotations (binomial test 95% confidence interval error bars). All gene expression data is from CAGE (Methods). (D-F) Histograms of (D) GC content, (E) conservation (PhyloP score) (Siepel et al., 2005) and (F) Hi-C compartment eigenvalues in IMR90. Positive PhyloP scores indicate evolutionary conservation, positive scores indicate accelerated evolution, and scores near zero indicate neutral evolution. Hi-C compartment values are computed according to the method of Lieberman-Aiden et al. (2009) (Methods). Positive values indicate open compartment, negative values indicate closed compartment. (G) Enrichment of each domain type with respect to IMR90 replication time (early vs. late) and replication time dynamics across cell types (constitutive vs. switching) (V Dileep, F Ay, J Sima, WS Noble, DM Gilbert et al., unpublished data) (Methods). (H) CTCF motif density for each domain type. Dashed line indicates genome-wide average. (I) Fraction of CTCF motifs bound (overlapping a CTCF peak) in IMR90 (binomial test 95% confidence interval error bars). Dashed line indicates average over all motifs.

The second known type of repressive domain is best known as “facultative heterochromatin” but is also sometimes referred to as BLOCs or Polycomb-repressed chromatin (Morey and Helin, 2010; Pauler et al., 2009). Facultative heterochromatin is regulated by the Polycomb complex and is associated with the histone modification H3K27me3. Facultative heterochromatin is thought to repress tissue-specific genes in cells where they are inactive. As expected, one output domain type “FAC” has all the known properties of facultative heterochromatin. FAC domains are marked by H3K27me3 (Figure 2A,D), and they are enriched for genes (Figure 3A), GC content (Figure 3D) and conservation (Figure 3E), but strongly depleted for gene expression relative to an average across cell types (Figure 3B), indicating that FAC domains have a direct repressive effect. FAC domains are mixed between the open and closed compartments, indicating that facultative repression is independent of compartment-driven repression (Figure 3F). However, FAC domains are almost completely absent from the annotation of the embryonic stem cell line H1-hESC, consistent with previous observations that H3K27me3 does not form domains in embryonic stem cells but rather occurs only at so-called poised or bivalent promoters (Supplementary Figure 4) (Bernstein et al., 2006).

Other semi-automated genome annotation analyses have reported a third type of repressive domain, characterized by a lack of signal from any mark, termed “quiescent domains” (Hoffman et al., 2012; Ernst and Kellis, 2010; Filion et al., 2010; Julienne et al., 2013). We identified this domain type as the QUI label (Figure 2A). Note that Segway marginalizes over missing data rather than setting the values to zero (Supplementary Note 3), so the QUI label is not simply an artifact of unmappable regions. QUI domains are highly depleted for genes (Figure 3A) and occur in the closed compartment (Figure 3F). QUI domains are depleted for transcription factor motifs but, unlike FAC and CON domains, are not depleted for transcription factor binding at motifs, indicating that QUI chromatin does not have a direct repressive effect (Figure 3H,I). The mechanism behind the activity of QUI domains is unknown, but these results are consistent with a model in which QUI domains lack any activating signals but are not directly repressed.

### Active domains are divided between broad and specific gene expression

Previous studies of human domains have focused on various types of repressive domains but have assigned all active chromatin to one domain category (Julienne et al., 2013; Wen et al., 2009; Pauler et al., 2009). However, studies in other organisms have reported multiple types of active domains (Filion et al., 2010; Liu et al., 2011). We therefore investigated whether our IMR90 annotation can be used to identify types of human active domains. We found that active domains in IMR90 can be split into BRD ("broad expression") domains, characterized by transcription-associated marks such as H3K36me3, and SPC ("specific expression") domains, characterized by regulatory marks such as H3K27ac. Both domain types are highly enriched for genes (Figure 3A). However, while genes in BRD domains are mostly expressed across all cell types, a much larger fraction of active genes in SPC domains are expressed only in a small number of cell types (Figure 3C). Furthermore, when a gene is in a SPC domain, that gene is expressed at a much higher level than that gene’s average across cell types, suggesting that SPC domains are highly activating (Figure 3B). In contrast, while genes in BRD domains are highly expressed, this high expression generally occurs consistently across cell types, indicating that BRD domains do not necessarily directly promote expression (Figure 3B). Moreover, while both BRD and SPC domains are generally early-replicating in IMR90, regions covered by SPC domains typically switch replication time between cell types, while regions covered by BRD domains are typically early replicating in all cell types (Figure 3G). These results suggest a model in which genes performing housekeeping functions such as DNA repair have strong promoters but little other regulation, whereas genes specific to a given tissue are regulated by a complex web of regulatory elements, allowing the genome to specify precise conditions under which the gene is active.

To test this hypothesis, we computed the enrichment of Gene Ontology (GO) terms for genes in BRD and SPC domains respectively (Gene Ontology Consortium, 2000; Boyle et al., 2004). We found that genes in BRD domains were enriched for housekeeping functions such as cell cycle and DNA repair, while genes in SPC domains were enriched for IMR90-specific developmental functions such as vasculature development and stimulus response (Supplementary Tables 3–4, Supplementary Figure 5). In order to avoid hindsight bias, before looking at these GO term enrichments, we mixed the enriched terms with an equal number of decoy terms matched according to the number of genes associated with each term, and manually labeled which terms matched our hypothesized functions for each domain (housekeeping for BRD, IMR90-specific for SPC). We correctly identified 21/32 BRD enrichments (*p* = 0.055) and 54/64 SPC enrichments (*p* = 1.4 × 10^-6^). This demonstrates that active regions can be divided into domains of broadly-expressed housekeeping genes and domains of specifically-expressed developmental genes. To our knowledge, this is the first time a split between domains of broad and specific expression has been reported in human cells.

### Lamina association is driven by a complex structure of domains

Previous work has shown that some repressive domains are marked with the histone modification H3K9me2, associate with the factor lamin B1 and localize to the nuclear lamina (Guelen et al., 2008; Wen et al., 2009). We found that comparing lamina association to domain annotations based on many data sets reveals a much more complex interaction than comparing to each mark individually (Figure 4). As expected, repressive domains (QUI and FAC) are enriched inside lamina-associating chromatin domains, while active domains are depleted. However, this analysis also reveals that CON domains are depleted immediately inside lamina-associating domain boundaries while being comparatively enriched at their centers. In contrast, FAC domains are highly enriched at lamina-associating domain boundaries while being comparatively depleted at their centers. In addition, while active domains (SPC and BRD) are depleted inside lamina-associating domains, they are highly enriched directly outside their boundaries. These observations suggest that lamina-associating domains form around a core of repressed chromatin and spread until they hit a strong active element.

**Figure 4:**
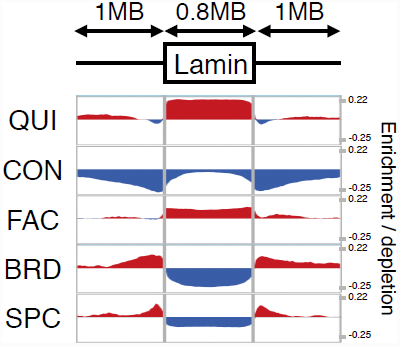
Enrichment of each domain type with respect to lamina-associating domain boundaries. X axis indicates position with respect to lamina-associating domains, with each domain stretched or shortened to the median length of 0.8 MB. Y axis indicates label enrichment or depletion (log(obs/expected)).

### Developmentally-consistent domain boundaries are marked by identifiable sequence elements

Previous research has shown that domain boundaries tend to be consistent between cell types even when the state of the domain changes. For example, when a region’s replication time is perturbed by leukemia, the boundaries of the resulting replication domain tend to occur at the same positions as developmental replication timing domain boundaries (Ryba et al., 2012). However, the cause of these consistent domain boundaries remains unclear. We investigated the consistency of domain boundaries using our domain annotations. As expected, domain boundaries frequently occurred at consistent positions across cell types, even when the domains’ state changed (Figure 5A). To identify these consistent domain boundaries, we combined all boundaries occurring in at least one cell type and merged boundaries within 50 kb. We defined groups of five or more boundaries as *consistent* (Methods) (Figure 5B). As expected, these consistent boundaries are enriched for replication domain boundaries, but many consistent domain boundaries do not overlap a replication domain boundary (Supplementary Figure 6). We additionally found that consistent domain boundaries are highly enriched for promoters and CTCF motifs, suggesting that these elements may drive domain boundary formation (Figure 5C,D).

**Figure 5:**
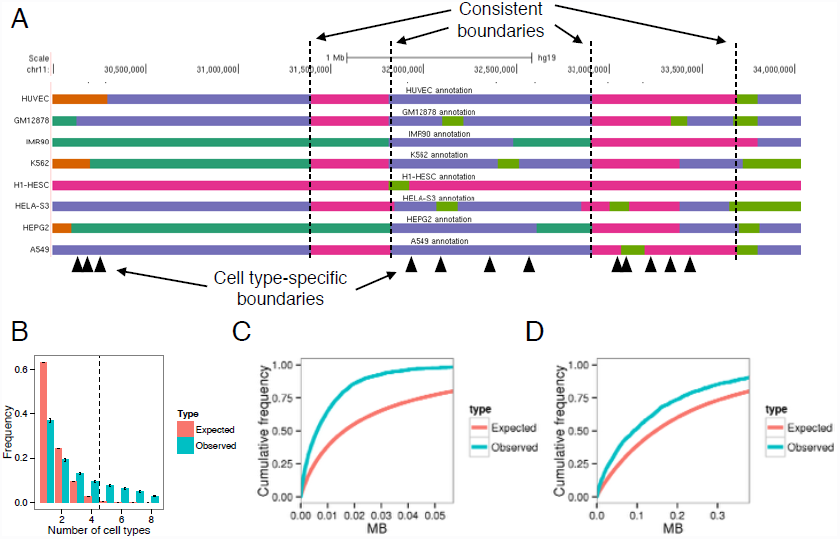
(A) Example of consistent domains. (B) Distribution of number of overlapping boundaries compared to a permutation control (binomial test 95% confidence interval error bars). Vertical dashed line denotes consistent boundary threshold. (C,D) Cumulative density of distances from consistent domain boundaries to the nearest (C) promoter and (D) distal CTCF motif. Distal CTCF motifs are defined as all CTCF motifs more than 5 kb from a promoter.

### Using GBR to transfer information between cell types improves accuracy of predicting functional elements

Graph-based regularization can also be used for the seemingly-unrelated task of transferring information from well-studied cell types for the annotation of cell types with limited available data (Figure 6A). Existing SAGA methods work well on data from a single cell type, but integrating information between cell types remains an open problem. Existing methods for using data from multiple cell types for genome annotation fail to effectively address this problem (Supplementary Note 4). We propose a novel strategy for leveraging information from well-studied cell types using the pairwise prior that if two positions received the same label in many well-studied cell types, then they should be more likely to receive the same label in the target cell type (Figure 6A). To express this pairwise prior, we first perform a Segway annotation (without GBR) of each well-studied cell type and create a GBR graph which connects each pair of positions with weight proportional to the number of cell types in which the pair receive the same label, placing higher weight on cell types similar to the cell type of interest (Methods). We then use this graph in combination with the data sets available in the target cell type to produce an annotation of this cell type. Note that this GBR graph represents an entirely different type of information from the graphs used to represent Hi-C data in the previous sections, despite the fact that both types of data are represented as a graph.

**Figure 6:**
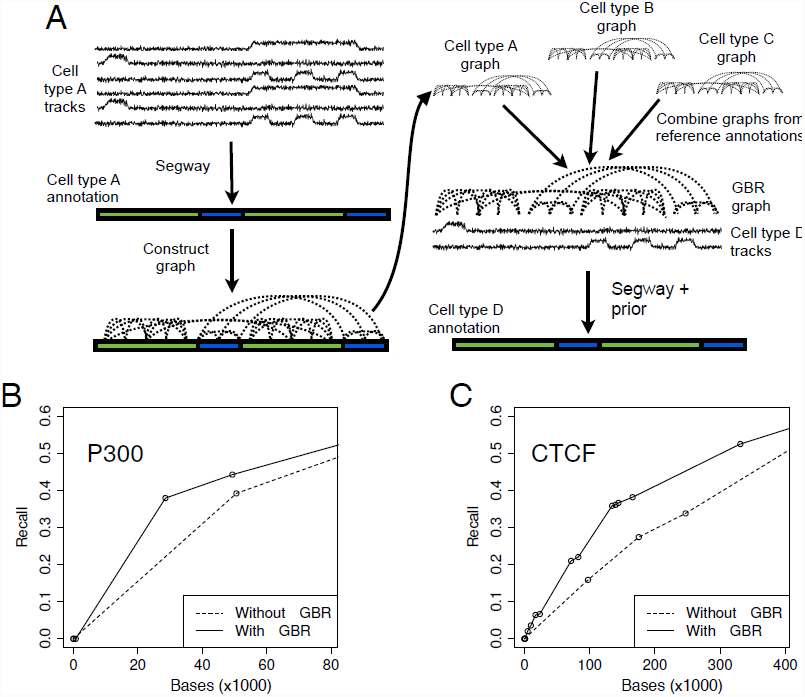
(A) Strategy for using GBR to transfer information between cell types. (B,C) Performance of predicting the locations of GM12878 (B) p300 and (C) CTCF ChIP-seq peaks with and without using GBR to integrate information from K562. Each plot shows the fraction of elements detected as a function of the number of bases predicted (Methods). Results are shown on the test set (10 Mbp). Model training and label ordering was performed on the training set (10 Mbp). The X axis is plotted up to 1 kbp times the total number of elements. These plots can be interpreted, for example, in the context of an enhancer validation experiment, in which case it shows how many sequences would need to be tested in order to discover a certain number of enhancers.

To demonstrate the efficacy of this approach, we evaluated whether GBR improves an annotation’s ability to predict enhancers and insulators. We simulated the case where the lymphoblastoid cell type GM12878 has only eight histone modifications available, a panel of data types similar to that assayed by Roadmap Epigenomics on hundreds of human tissues (Supplementary Note 2). Because there are enough well-studied cell types to ensure that at least one reference is reasonably closely related to any cell type of interest, we used the related leukemia cell type K562 as reference. We annotated GM12878 using these eight histone modifications and a GBR graph derived from an annotation of K562 (Methods). Incorporating information from K562 this way greatly improved the accuracy with which the annotation detected enhancers and insulators (Figure 6B,C). We evaluated the performance with which the GM12878 annotation predicts a certain type of functional element by ordering the labels by their enrichment for the element on a training set and evaluating the recall as more labels are added (Methods). The GBR annotation detects one third of p300 binding sites (a proxy for enhancers (Visel et al., 2009)) by predicting just 25 kb as p300-binding, while the annotation produced without GBR predicts 43 kb before it detects this many sites (Figure 6B). Likewise, the GBR annotation detects one third of CTCF binding sites (a proxy for insulators (Burgess-Beusse et al., 2002)) by predicting 124 kb, compared to 241 kb without GBR (Figure 6C). Because the algorithm was not given any knowledge of enhancers or insulators as input, it is reasonable to expect that the annotations achieve similar performance at detecting other types of functional elements, for which we do not have gold-standard examples and therefore cannot evaluate against. These results demonstrate that GBR effectively leverages information from a reference cell type and therefore provides a method for producing high-quality annotations of the hundreds of cell types with limited available data.

## Discussion

We introduced graph-based regularization (GBR), a method which allows probabilistic models to integrate a pairwise prior while maintaining efficient inference. We used GBR to model chromatin conformation data and thereby jointly model all available data types for the study of chromatin domains. To our knowledge, this represents the first method for integrating chromatin conformation information into SAGA methods without resorting to simplifying transformations. We showed that modeling Hi-C data with GBR improved the annotation’s ability to predict replication time and topological domains. In addition, because graph-based regularization is a general method, it will likely prove useful for other applications involving dynamic Bayesian networks, such as methods for locating genes or predicting copy number.

The ability to integrate Hi-C data into an annotation allowed us to study the relationship between types of domains by integrating all available data into a single annotation (Figure 7A). This analysis revealed a set of five domain types that encompass all previously-described domain types: (1) quiescent domains, which lack any activity, (2) constitutive heterochromatin, which represses permanently silent regions and is marked with the histone modification H3K9me3, (3) facultative heterochromatin, which represses cell type-specific regions and is marked with the histone modification H3K27me3, (4) broadly-expressed domains, which cover genes that are highly expressed in all cell types, and (5) specifically-expressed domains, which exhibit high regulatory activity and cover genes that are expressed in a small number of cell types.

**Figure 7:**
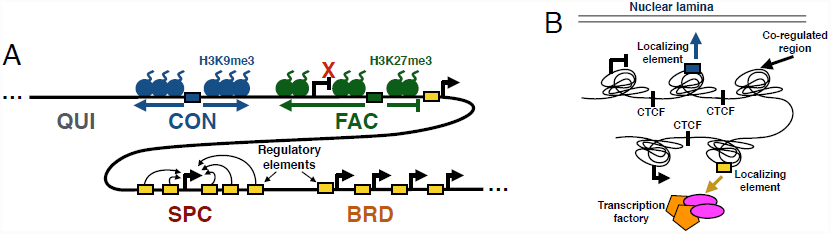
Models of (A) domain types and (B) co-regulated regions.

To our knowledge, domains of specific expression (SPC) have not been identified previously in human cells. These domains are likely the result of complex regulatory programs designed to precisely control the condition and level of genes important for a certain cell state or function. SPC domains are similar in some ways to dense clusters of regulatory elements important for cell identity known as superenhancers (Whyte et al., 2013; Lovén et al., 2013). However, there are only small number of known superenhancers (~300) and each is much smaller than a SPC domain (~10 kb compared to ~200 kb). Therefore, SPC domains and superenhancers may result from similar mechanisms, but on very different scales. However, the mechanisms underlying of both types of regions must be studied further in order to understand this relationship.

One likely mechanism of domain formation involves the spreading of heterochromatin (Weiler and Wakimoto, 1995; Talbert and Henikoff, 2006). Under this hypothesis, heterochromatin nucleates at silencing elements such as telomeres, repeats or repressed promoters and sequentially assembles along chromatin. Spreading heterochromatin has been demonstrated mechanistically in *Saccharomyces* and *Drosophila* and for the SIR, HP1 and Polycomb complexes. While SIR is unique to yeast, HP1 and Polycomb have orthologs in humans that drive constitutive and facultative heterochromatin, respectively. Under the spreading hypothesis, heterchromatin can be halted by the presence of a strong active element. This halting mechanism is consistent with the observation that active domains (especially SPC) are strongly enriched directly outside of lamina-associating domains.

The consistency of domain boundaries between cell types also suggests a model in which core regions are regulated as a unit (Phillips and Corces, 2009; Dixon et al., 2012) (Figure 7B). Under this hypothesis, these units self-interact as topological domains and are co-regulated through availability of regulatory factors and elements such as enhancers. These co-regulated units are thought to be delimited by localizing sequences, particularly CTCF sites. Under this model, each of our annotated domains is actually composed of several such neighboring co-regulated regions with the same state. Therefore, while profiling a small number of cell types has allowed us to define a small number of consistent domain boundaries, profiling more cell types may lead to a complete catalogue of potential boundary sites.

We have described five domain types because we this model allowed us to concisely summarize domain regulation, but we do not claim that this represents the “true” number of domain types. It is likely that new domain subtypes will be discovered in the future, thus increasing the number of known domain types. In addition, methods that discover the optimal number of domain types or that allow mixtures of domain types are an interesting direction for future work. To our knowledge, all existing SAGA methods require either a fixed number of labels (Day et al., 2007; Hoffman et al., 2012; Ernst and Kellis, 2010; Thurman et al., 2007; Lian et al., 2008; Filion et al., 2010) or a hyperparameter that indirectly controls the number of labels (Ho et al., 2014). A method that allows for “mixed” domain labels at a given process could potentially circumvent the manual merging process that we used to reduce an eight-label model to a five-label one.

Finally, we presented a method for transferring information from well-studied cell types using GBR in order to improve the quality and interpretability of annotations of cell types with limited available data. This method enables a new strategy for understanding cell types, in which a small number of assays are performed on each cell type of interest to determine the unique characteristics of this cell type, then Segway with GBR is used to combine this data with the large body of available information from well-studied cell types. This method has the additional benefit of matching the label semantics of the target cell types to the semantics of the reference annotations, which allows the label interpretation process to be performed automatically Because consortia such as ENCODE and Roadmap Epigenomics are already analyzing a large number of cell types with a small number of assays each, this strategy is immediately applicable. Determining which assays are most informative as input to this strategy is an interesting question for future work.

## Methods

### Histone modification, open chromatin and replication timing signal data

We acquired histone ChIP-seq, DNase-seq, and FAIRE-seq data for A549, K562, H1-hESC, GM12878, HeLa-S3, HepG2 and HUVEC from ENCODE and for IMR90 from Roadmap Epigenomics (ENCODE Project Consortium, 2012; Bernstein et al., 2010). We used a uniform signal-processing pipeline to generate a genome-wide vector for each data set, as described in (Hoffman et al., 2013). We also acquired Repli-seq data for IMR90 from ENCODE and smoothed this data using wavelet smoothing as described in (Thurman et al., 2007). We applied the inverse hyperbolic sine transform asinh 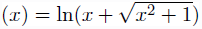 to all signal data. This transform is similar to the log transform in that it depresses the magnitude of extremely large values, but it is defined at zero and amplifies the magnitude of small values less severely than the log transform does. This transform has been shown to be important for reducing the effect of large values in analysis of genomics data sets (Johnson, 1949; Hoffman et al., 2012).

We acquired transcription factor ChIP-seq data from ENCODE. Peaks were called for each factor using MACS using an IDR threshold of 0.05 (Zhang et al., 2008; Landt et al., 2012).

We acquired CAGE expression data for 33 cell types from GENCODE (Harrow et al., 2012).

The full list of data sets used is available in Supplementary Table 1.

### Hi-C data

We used publicly available Hi-C data sets for two human cell lines (IMR90 and H1-hESC) (Dixon et al., 2012). We processed raw paired-end libraries with a pipeline that combines reads from two replicates per cell line, maps these reads, extracts the read pairs for which each end maps uniquely, and removes potential PCR duplicates. We then partitioned the human genome into a collection of non-overlapping 10 kb windows and assigned each end of a read pair to the nearest 10 kb window mid-point. This process yielded a 303,641×303,641 whole genome contact map. These contact maps consisted of both intra and interchromosomal contacts and contained only ~0.3% non-zero entries. We assigned statistical confidence estimates to these contact counts using the method Fit-Hi-C, which jointly models the random polymer looping effect and technical biases (Ay et al., 2014a). First, we applied the bias correction method ICE to the contact map to estimate a bias associated with each 10 kb locus, after eliminating all loci that have less than 50% uniquely mappable bases (Imakaev et al., 2012). Second, using these computed biases and raw contact maps as input, we estimated a *p-*value of interaction for each pair of 10 kb loci with non-zero contact counts *(p-*value was set to 1 for pairs with zero contacts). We used a slightly modified version of the original Fit-Hi-C algorithm which handles inter- as well as intrachromosomal contacts and omits the refinement step for fast computation. Because Fit-Hi-C normalizes for 1D genomic distance, the majority of significant contacts were at long distances (Supplementary Figure 7). Note that while the data sets we used have insufficient coverage to identify many high-confidence contacts at 10kb resolution, Segway with GBR aggregates information over roughly 400kb in order to make each domain call, so individual high-confidence interactions are not necessary.

We computed the genome chromatin compartment using eigenvalue decomposition on the normalized contact maps of IMR90 and H1-ESC cell lines at 1 Mb resolution as described in (Lieberman-Aiden et al., 2009). For each chromosome, we calculated the Pearson correlation between each pair of rows of the intrachromosomal contact matrix and applied eigenvalue decomposition to the correlation matrix. Similar to (Lieberman-Aiden et al., 2009), we used the second eigenvector in cases where the first eigenvector values were either all positive or all negative to define the compartments. We used average GC content to map signs of eigenvectors to either open (higher GC content) or closed chromatin compartments.

### Graph-based regularization

In a SAGA method, we are given a set of vertices *V* that index a set of *n* = |*V*| random variables *X*_*V*_ = {*X*_1_,...,*X*_*n*_} and a probability distribution parameterized by θ, *p*_*θ*_(*X*_*H*_,*X*_*O*_). Different SAGA methods employ different distributions *P*_*θ*_. Graph-based regularization could be applied to any probabilistic model, but in this work we use the Segway model (Supplementary Methods) because it can handle real-valued and missing data, and it can use non-geometric segment length distributions. We denote random variables with capital letters (e.g., *X*_*H*_) and instantiations of variables with lower-case (e.g., *x*_*H*_ ∈ domain(*X*_*H*_)). We use capitals to denote sets and lowercase to denote values (e.g., *X*_*h*_ for *h* ∈ *H*).

Training the model involves a set of observed data 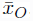, where a subset of variables *O* ⊆ *V* is observed, and the remainder *H* = *V \ O* are hidden. The maximum likelihood training procedure optimizes the objective

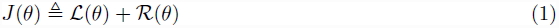

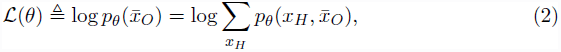

where 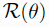 is a regularizer that expresses prior knowledge about the parameters. Many regularizers are used in practice, such as the *𝓁*_2_ or *𝓁*_1_ norms, which encourage parameters to be small or sparse respectively.

Dynamic programming algorithms such as the forward-backward algorithm can be used to perform inference in SAGA models, because all such existing models have dependencies in the form of a chain. That is, the variables associated with position *i* depend only on the variables associated with positions *i –* 1 and *i* + 1. Examples of such chain-structured models include hidden Markov models and dynamic Bayesian networks. However, these dynamic programming algorithms do not apply if a pairwise prior is added to the model, since the prior may have an arbitrary structure. Several techniques have been proposed to handle models with arbitrary structure (Supplementary Note 5). However, none of these techniques are optimal for expressing a pairwise prior.

Therefore, we instead employ a novel strategy based on *posterior regularization* (Ganchev et al., 2010) to integrate this prior. This is done by introducing an auxiliary joint distribution *q*(*X*_*H*_), placing a regularizer on *q*(*X*_*H*_), and encouraging *q* to be similar to *p*_*θ*_ through a KL divergence penalty. The regularizer is

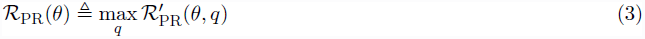

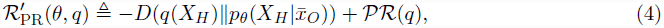

where *D*(⋅||⋅) is the KL divergence *D*(*p*(*X*_*H*_)||*q*(*X*_*H*_)) = *∑*_*X*_*H*__ *p*(*x*_*H*_) log(*p*(*x*_*H*_)/*q*(*x*_*H*_)) and 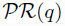 is a posterior regularizer that expresses prior knowledge about the posterior distribution. KL divergence measures the dissimilarity of probability distributions, such that *D*(*p*||*q*) is zero if the distributions are identical and can be arbitrarily large if they are not. Several posterior regularizers have been proposed in the past, such as those that require posteriors to satisfy constraints in expectation (Ganchev et al., 2010).

We propose a new type of posterior regularizer that expresses a pairwise prior (Figure 8). We are given a weighted, undirected regularization graph over the hidden variables *G*_*R*_ = (*H, E*_*R*_), where *E*_*R*_ ⊆ *H* x *H* is a set of edges with non-negative similarity weights 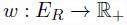, such that a large *w*(*u, v*) indicates that we have strong belief that *X*_*u*_ and *X*_*v*_ should be similar. (We describe how we generate this graph in the next two sections.) For a distribution *p*(*X*_*H*_), let 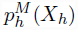 indicate the marginal distribution over 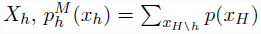. Let *λ*_*G*_ be a hyperparameter controlling the strength of regularization. The posterior regularizer is

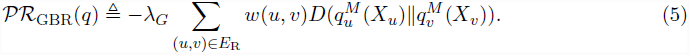

**Figure 8:**
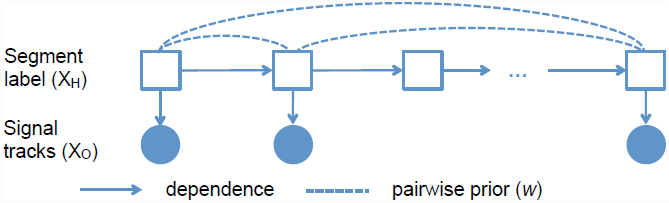
GBR model. Squares and circles denote discrete and continuous random variables respectively. Filled-in and unfilled shapes denote observed and unobserved variables, respectively.

Thus the full objective is

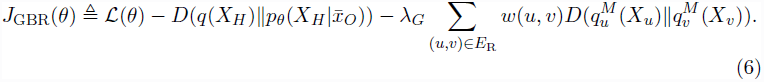

We term this strategy of adding graph-based penalties *graph-based regularization* (GBR).

### GBR optimization

We have developed a novel algorithm for efficiently optimizing *J*_GBR_ in *q*. This algorithm alternates between using a method for probabilistic inference such as the forward-backward algorithm and applying a message passing algorithm over the regularization graph *G*_*R*_. In the inference step, the model receives evidence from the message passing step in the form of a “virtual evidence” distribution 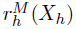 over each variable *h*. These virtual evidence distributions are used in conjunction with the original SAGA model to compute a posterior distribution over the labels using any algorithm for probabilistic inference on dynamic Bayesian networks, such as belief propagation or the forward-backward algorithm.

In the message passing step, the algorithm updates *r*^*M*^ to minimize the Kullback-Leibler penalties in the objective function *J*_GBR_. This message passing step is itself performed using an alternating optimization algorithm, which passes messages over the regularization graph *G*_*R*_. This algorithm is similar to one originally developed for the field of semi-supervised learning (Subramanya and Bilmes, 2011).

The inference and message passing steps are iterated until convergence. These two updates are linear in the number of variables (for chain-structured models, which include all existing SAGA methods) and linear in the degree of the regularization graph, respectively. The algorithm exhibits monotonic convergence, similar to the EM algorithm. We derive the algorithm for optimizing *J*_GBR_ and prove its convergence in Supplementary Note 1.

### GBR graph for incorporating Hi-C data

When we are using GBR to incorporate Hi-C data, we are given a matrix of contact *p*-values 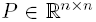, generated from a matrix of contact counts as described above. To remove noise and decrease the degree of the graph, we removed all contacts with uncorrected p-value *p >* 10^-6^ and multiplied the remaining *p-*values by 10^6^, similar to a Bonferroni correction. Note that due to the large number of hypotheses, performing a full Bonferroni correction would result in very few contacts. Moreover, the graph weights allow the algorithm to take into account the strength of each connection, so the choice of 10^6^ was made for computational, not statistical, reasons. We computed the weights as

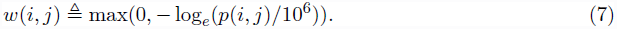

As with the graph for transferring information between cell types, the multiplicative scale of the weights is arbitrary, since it is controlled by the graph weight hyperparameter λ_*G*_. We used only intrachromosomal contacts for forming the GBR graph. To produce a GBR graph representing cell type-consistent chromatin conformation used in the domain annotation of eight cell types, we added the edge weights from the IMR90 and H1-hESC Hi-C GBR graphs.

### GBR graph for annotation of multiple cell types

When we are using GBR to transfer information about cell type *A* to improve annotation of cell type *B*, we are given an annotation 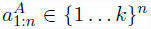 of cell type *A*, produced without GBR. We construct a GBR graph from this annotation by connecting each pair of positions that received the same label in *a*^*A*^ with an undirected edge of weight 1. Note that the weight is arbitrary, since it is scaled by the regularization parameter λ_*G*_. To mitigate the problem of quadratic growth in the degree of this graph, we randomly subsampled this graph such that each node had outgoing degree 17 ≈ log_*e*_(n). We chose this graph degree because a randomly-subsampled graph with *n* log_*e*_ *n* edges has the same connected components as the full graph with high likelihood (Erdős and Rényi, 1960), and our experiments on synthetic data (not shown) showed that the sparse graph performed similarly to a complete graph.

### Circular permutation

As a null model for several experiments, we performed a circular permutation of the genome along each chromosome arm as follows. We randomly choose a translation fraction *θ* ∈ [0,1]. For each coordinate *i* ∈ {1... n} within a chromosome arm that spans the range [a, b), we translate *i* to *t*(*i*), where

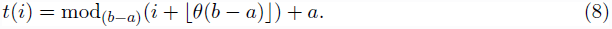

To circularly permute a genome feature, such as an annotation or a Hi-C contact map, we translate each element from position *i* to *t*(*i*). Thus, when a circularly-permuted feature is compared to an un-permuted feature, all positional correspondence between permuted and un-permuted features are removed, but each feature’s spatial patterns are preserved. In each case, we performed this permutation 200 times and report the average over all permutations. If the feature includes any centromere- or telomere-defined elements, we remove these as a preprocessing step.

### Topological domain agreement

To evaluate the degree to which an annotation *A* matches a set of topological domains, we computed the number *c*_*d*,*𝓁*_ of bases by which domain *d* is covered by label *𝓁.* We then computed 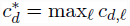 to be the number of bases covered by the highest-coverage label for domain *d*, and divided 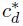 by the length of *d* to produce 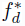, the fraction of *d* covered by its plurality label. The agreement 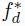 takes its maximum value of 1 if the domain *d* is covered by exactly one annotation label. We computed the raw genome-wide agreement 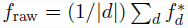.

This raw genome-wide agreement *f*_raw_ can be improved simply by increasing the length of segments and decreasing the number of labels. Therefore, we circularly permuted *A* to form *A*^*p*^, and used this permuted annotation to compute 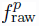 Finally, we computed the *topological domain agreement* 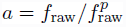 as the ratio of unpermuted and permuted raw agreements. This normalized agreement is large when the annotation has small segments that exactly match the topological domains and is small when the annotation’s segments are not correlated with topological domains.

### Signal variance explained

To evaluate the similarity between a genome-wide signal vector and a genome annotation, we use the following measure, which we term the *variance explained.* We are given a genome annotation with *k* labels a_1:*n*_ ∈ {1... *k*}^*n*^ and a vector 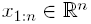. We compute the signal mean over the positions assigned a given label *𝓁* as

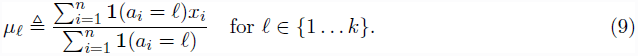

We define a predicted signal vector 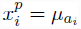. and compute the the prediction error as 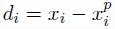. We compute the residual standard deviation of the signal vector as

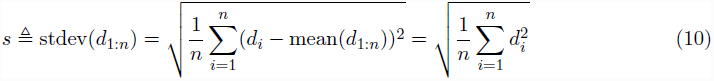

The last equality holds because mean(d_1:*n*_) = 0 by construction. We define the *variance explained* (VE) for annotation *a* and signal vector *x* as VE ≜ stdev(*d*_1:*n*_) – stdev(*x*_1:*n*_). VE is bounded by the range [0, stdev(*x*_1:*n*_)]. VE is a measure of the extent to which a genome-wide signal data set and annotation are similar, where higher values indicate better agreement.

### Genomic element prediction

We form a classifier for a set of genomic elements based on an annotation using the following strategy. We are given a genome annotation with *k* labels *α*_1:*n*_ ∈ {1... *k*}^*n*^ and a set of positions *S* ⊆ {1... *n*} which represent some set of elements of interest, such as enhancers or CTCF binding sites. Define *A*_*l*_ = { *i* | *a*_*i*_ = *ℓ* } to be the positions annotated by label *ℓ* To avoid biases caused by differing-size elements, we assume that each element occupies just 1 bp. In the case of larger elements (such as MACS-called TF binding sites, which are ~200 bp), we define each element as the middle base pair of the range.

For each label, we compute the predictive precision of label *ℓ* as

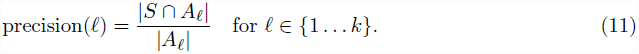

We rank the labels in decreasing order of their precision on a training set to get an order *s*_1:*k*_∈ {1...*k*}^*k*^.

Using this ordering, we form *k* predictors, 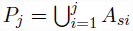 for *j* ∈ {1... *k*}. The true positives and false positives of a predictor *P* are TP(*P*) = *P* ∩ *S* and FP(P) = *P*∩({1...*n*}\*S*) respectively. The predictors are in order of decreasing stringency—that is, *P*_*j–*1_ ⊆ *P*_*j*_.

We can trace out the full sensitivity-specificity tradeoff (such as for an ROC or PR curve), by interpolating between each successive pair of predictors. To interpolate between a pair of predictors *P*_*j*_ and *P*_*j*+1_, we form an interpolated predictor *P*_*j,j+*1,*θ*_ by sampling each position *i* ∈ *Pj\P*_*j*–1_ with probability *θ* ∈ [0,1]. The expected number of true positives and false positives of an interpolated predictor *P*_*j,j+*1,*θ*_ can be shown to be

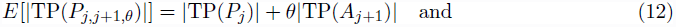

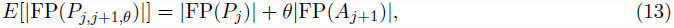

respectively. We report our performance using a test set disjoint from the training set used to order the labels.

### Developmental replication domains

In order to evaluate the replication timing dynamics of different types of domains, we used a four-label (constitutive early/late, switching early/late) annotation of the human genome using published replication timing data for 16 different human cell types, gathered by V Dileep, F Ay, J Sima, WS Noble, DM Gilbert et al. (unpublished). This annotation first windowed replication timing data into 40 kb bins and then determined for each window whether it replicates early (RT value > 0.5) or late (RT value < −0.2) in all cell types. Such windows with consistent timing profiles across all cell types were labeled as “constitutively early” and “constitutively late”, respectively. The remaining windows were labeled as either switching or left unlabeled. Switching windows are determined as those with an absolute value of replication timing larger than 0.5 in all cell types but with an opposite sign than others in at least one cell type. Switching windows that are early and late replicating in IMR90 were labeled as switching early and switching late, respectively.

### Consistent domain boundaries

When we annotated domains in eight cell types, we found that domain boundaries were shared between annotations much more often than would be expected by chance. To identify developmentally-consistent domain boundaries, we first formed a list of all segment boundaries that occurred in at least one cell type. For each boundary, we computed the number of cell types with boundaries within 50 kb. We formed a set of representative by greedily selecting the boundary with the most nearby boundaries as a representative, removing all boundaries near the representative from the list, and repeating the process until no two boundaries in the list were within 50 kb of one another. While this problem is an instance of the NP-hard set cover problem, the greedy approach is guaranteed to result in a constant-factor approximation of optimal (Nemhauser et al., 1978). This yielded a set of 13,906 boundary groups, each more than 50 kb from all other groups. We defined the 2,967 boundary groups composed of at least five boundaries as consistent boundaries.

## Data access

Domain annotations and code for Segway with GBR is available online at http://noble.gs.washington.edu/proj/gbr.

## References

Akhtar W, de Jong J, Pindyurin AV, Pagie L, Meuleman W, de Ridder J, Berns A, Wessels LF, van Lohuizen M, and van Steensel B. 2013. Chromatin position effects assayed by thousands of reporters integrated in parallel. Cell 154: 914–927.

Altun Y, Belkin M, and Mcallester DA. 2005. Maximum margin semi-supervised learning for structured variables. In Advances in Neural Information Processing Systems, pp. 33–40.

Ay F, Bailey TL, and Noble WS. 2014a. Statistical confidence estimation for Hi-C data reveals regulatory chromatin contacts. Genome Research 24: 999–1011.

Ay F, Bunnik EM, Varoquaux N, Bol SM, Prudhomme J, Vert JP, Noble WS, and Le Roch KG. 2014b. Three-dimensional modeling of the *P. falciparum* genome during the erythrocytic cycle reveals a strong connection between genome architecture and gene expression. Genome Research 24: 974–988.

Bernstein BE, Mikkelsen TS, Xie X, Kamal M, Huebert DJ, Cuff J, Fry B, Meissner A, Wernig M, Plath K, et al.. 2006. A bivalent chromatin structure marks key developmental genes in embryonic stem cells. Cell 125: 315–326.

Bernstein BE, Stamatoyannopoulos JA, Costello JF, Ren B, Milosavljevic A, Meissner A, Kellis M, Marra MA, Beaudet AL, Ecker JR, et al.. 2010. The NIH Roadmap Epigenomics Mapping Consortium. Nature Biotechnology 28: 1045–1048.

Bickmore WA and van Steensel B. 2013. Genome architecture: Domain organization of interphase chromosomes. Cell 152: 1270–1284.

Biesinger J, Wang Y, and Xie X. 2013. Discovering and mapping chromatin states using a tree hidden Markov model. Bioinformatics 14: S4.

Bilmes J and Zweig G. 2002. The Graphical Models Toolkit: An open source software system for speech and time-series processing. In Proceedings of the IEEE International Conference on Acoustics, Speech, and Signal Processing.

Bishop C. 1995. Neural Networks for Pattern Recognition. Oxford UP, Oxford, UK.

Boyle EI, Weng S, Gollub J, Jin H, Botstein D, Cherry JM, and Sherlock G. 2004. Go::TermFinder – open source software for accessing Gene Ontology information and finding significantly enriched Gene Ontology terms associated with a list of genes. Bioinformatics 20: 3710–3715.

Burgess-Beusse B, Farrell C, Gaszner M, Litt M, Mutskov V, Recillas-Targa F, Simpson M, West A, and Felsenfeld G. 2002. The insulation of genes from external enhancers and silencing chromatin. Proceedings of the National Academy of Sciences of the United States of America 99: 16433.

Chakalova L, Debrand E, Mitchell JA, Osborne CS, and Fraser P. 2005. Replication and transcription: Shaping the landscape of the genome. Nature Reviews Genetics 6: 669–677.

Das D and Petrov S. 2011. Unsupervised part-of-speech tagging with bilingual graph-based projections. In NAACL, pp. 600–609.

Das D and Smith N. 2011. Semi-supervised framesemantic parsing for unknown predicates. In Association for Computational Linguistics.

Day N, Hemmaplardh A, Thurman RE, Stamatoyannopoulos JA, and Noble WS. 2007. Unsupervised segmentation of continuous genomic data. Bioinformatics 23: 1424–1426.

Dekker J. 2008. Gene regulation in the third dimension. Science 319: 1793–1794.

Dixon JR, Selvaraj S, Yue F, Kim A, Li Y, Shen Y, Hu M, Liu JS, and Ren B. 2012. Topological domains in mammalian genomes identified by analysis of chromatin interactions. Nature 485: 376–380.

ENCODE Project Consortium. 2012. An integrated encyclopedia of DNA elements in the human genome. Nature 489: 57–74.

Erdős P and Rényi A. 1960. On the evolution of random graphs. Magyar Tud. Akad. Mat. Kutató Int. Közl 5: 17–61.

Ernst J and Kellis M. 2010. Discovery and characterization of chromatin states for systematic annotation of the human genome. Nature Biotechnology 28: 817–825.

Filion GJ, van Bemmel JG, Braunschweig U, Talhout W, Kind J, Ward LD, Brugman W, de Castro IJ, Kerkhoven RM, Bussemaker HJ, et al.. 2010. Systematic protein location mapping reveals five principal chromatin types in *Drosophila* cells. Cell 143: 212–224.

Ganchev K,Graça J, Gillenwater J, and Taskar B. 2010. Posterior regularization for structured latent variable models. Journal of Machine Learning Research 11: 2001–2049.

Gene Ontology Consortium. 2000. Gene ontology: tool for the unification of biology. Nature Genetics 25: 25–9.

Guelen L, Pagie L, Brasset E, Meuleman W, Faza MB, Talhout W, Eussen BH, de Klein A, Wessels L, de Laat W, et al.. 2008. Domain organization of human chromosomes revealed by mapping of nuclear lamina interactions. Nature 453: 948–951.

Harrow J, Frankish A, Gonzalez JM, Tapanari E, Diekhans M, Kokocinski F, Aken BL, Barrell D, Zadissa A, Searle S, et al.. 2012. GENCODE: the reference human genome annotation for The ENCODE Project. Genome Research 22: 1760–1774.

He L, Gillenwater J, and Taskar B. 2013. Graph-based posterior regularization for semi-supervised structured prediction. In Seventeenth Conference on Computational Natural Language Learning.

Ho JW, Jung YL, Liu T, Alver BH, Lee S, Ikegami K, Sohn KA, Minoda A, Tolstorukov MY, Appert A, et al.. 2014. Comparative analysis of metazoan chromatin organization. Nature 512: 449–452.

Hoffman MM, Buske OJ, Wang J, Weng Z, Bilmes JA, and Noble WS. 2012. Unsupervised pattern discovery in human chromatin structure through genomic segmentation. Nature Methods 9: 473–476.

Hoffman MM, Ernst J, Wilder SP, Kundaje A, Harris RS, Libbrecht M, Giardine B, Ellenbogen PM, Bilmes JA, Birney E, et al.. 2013. Integrative annotation of chromatin elements from ENCODE data. Nucleic Acids Res 41: 827–41.

Imakaev M, Fudenberg G, McCord RP, Naumova N, Goloborodko A, Lajoie BR, Dekker J, and Mirny LA. 2012. Iterative correction of Hi-C data reveals hallmarks of chromosome organization. Nature Methods 9: 999–1003.

Johnson NL. 1949. Systems of frequency curves generated by methods of translation. Biometrika pp. 149–176.

Julienne H, Zoufir A, Audit B, and Arneodo A. 2013. Human genome replication proceeds through four chromatin states. PLoS Computational Biology 9: e1003233.

Kuhn HW. 1955. The Hungarian method for the assignment problem. Naval Research Logistics Quarterly 2: 83–97.

Lachner M, O’Sullivan RJ, and Jenuwein T. 2003. An epigenetic road map for histone lysine methylation. Journal of cell science 116: 2117–2124.

Landt SG, Marinov GK, Kundaje A, Kheradpour P, Pauli F, Batzoglou S, Bernstein BE, Bickel P, Brown JB, Cayting P, et al.. 2012. ChIP-seq guidelines and practices of the ENCODE and modENCODE consortia. Genome Research 22: 1813–1831.

Lian H, Thompson W, Thurman RE, Stamatoyannopoulos JA, Noble WS, and Lawrence C. 2008. Automated mapping of large-scale chromatin structure in ENCODE. Bioinformatics 24: 1911–1916.

Lieberman-Aiden E, van Berkum NL, Williams L, Imakaev M, Ragoczy T, Telling A, Amit I, Lajoie BR, Sabo PJ, Dorschner MO, et al.. 2009. Comprehensive mapping of long-range interactions reveals folding principles of the human genome. Science 326: 289–293.

Liu T, Rechtsteiner A, Egelhofer TA, Vielle A, Latorre I, Cheung MS, Ercan S, Ikegami K, Jensen M, Kolasinska-Zwierz P, et al.. 2011. Broad chromosomal domains of histone modification patterns in C. elegans. Genome Research 21: 227–236.

Lovén J, Hoke HA, Lin CY, Lau A, Orlando DA, Vakoc CR, Bradner JE, Lee TI, and Young RA. 2013. Selective inhibition of tumor oncogenes by disruption of super-enhancers. Cell 153: 320–334.

Misteli T. 2007. Beyond the sequence: Cellular organization of genome function. Cell 128: 787–800.

Morey L and Helin K. 2010. Polycomb group protein-mediated repression of transcription. Trends in biochemical sciences 35: 323–332.

Neal R and Hinton G. 1999. A view of the EM algorithm that justifies incremental, sparse, and other variants. In Learning in graphical models, pp. 355–368. MIT Press.

Nemhauser GL, Wolsey LA, and Fisher ML. 1978. An analysis of approximations for maximizing submodular set functionsi. Mathematical Programming 14: 265–294.

Pauler FM, Sloane MA, Huang R, Regha K, Koerner MV, Tamir I, Sommer A, Aszodi A, Jenuwein T, and Barlow DP. 2009. H3k27me3 forms blocs over silent genes and intergenic regions and specifies a histone banding pattern on a mouse autosomal chromosome. Genome Research 19: 221–233.

Phillips JE and Corces VG. 2009. CTCF: master weaver of the genome. Cell 137: 1194–1211.

Pope BD, Ryba T, Dileep V, Yue F, Wu W, Denas O, Vera DL, Wang Y, Hansen RS, Canfield TK, et al.. 2014. Topologically associating domains are stable units of replication-timing regulation. Nature 515: 402–405.

Ryba T, Battaglia D, Chang BH, Shirley JW, Buckley Q, Pope BD, Devidas M, Druker BJ, and Gilbert DM. 2012. Abnormal developmental control of replication-timing domains in pediatric acute lymphoblastic leukemia. Genome Research 22: 1833–1844.

Ryba T, Hiratani I, Lu J, Itoh M, Kulik M, Zhang J, Schulz TC, Robins AJ, Dalton S, and Gilbert DM. 2010. Evolutionarily conserved replication timing profiles predict long-range chromatin interactions and distinguish closely related cell types. Genome Research 20: 761–770.

Sheffield NC, Thurman RE, Song L, Safi A, Stamatoyannopoulos JA, Lenhard B, Crawford GE, and Furey TS. 2013. Patterns of regulatory activity across diverse human cell types predict tissue identity, transcription factor binding, and long-range interactions. Genome Research 23: 777–788.

Siepel A, Bejerano G, Pedersen JS, Hinrichs AS, Hou M, Rosenbloom K, Clawson H, Spieth J, Hillier LW, Richards S, et al.. 2005. Evolutionarily conserved elements in vertebrate, insect, worm, and yeast genomes. Genome Research 15: 1034–1050.

Subramanya A and Bilmes J. 2011. Semi-supervised learning with measure propagation. Journal of Machine Learning Research 12: 3311–3370.

Subramanya A, Petrov S, and Pereira F. 2010. Efficient graph-based semi-supervised learning of structured tagging models. In Proc. of EMLNP 2010, pp. 167–176. Association for Computational Linguistics.

Supek F, Bošnjak M, Škunca N, and Šmuc T. 2011. REVIGO summarizes and visualizes long lists of gene ontology terms. PloS ONE 6: e21800.

Talbert PB and Henikoff S. 2006. Spreading of silent chromatin: inaction at a distance. Nature Reviews Genetics 7: 793–803.

Thurman RE, Day N, Noble WS, and Stamatoyannopoulos JA. 2007. Identification of higher-order functional domains in the human ENCODE regions. Genome Research 17: 917–927.

Visel A, Blow MJ, Li Z, Zhang T, Akiyama JA, Holt A, Plajzer-Frick I, Shoukry M, Wright C, Chen F, et al.. 2009. ChIP-seq accurately predicts tissue-specific activity of enhancers. Nature 457: 854–858.

Wainwright M and Jordan M. 2008. Graphical models, exponential families, and variational inference. Foundations and Trends in Machine Learning 1: 1–305.

Warga J. 1963. Minimizing certain convex functions. Journal of the Society for Industrial and Applied Mathematics 11: 588–593.

Weiler KS and Wakimoto BT. 1995. Heterochromatin and gene expression in *Drosophila*. Annual review of genetics 29: 577–605.

Wen B, Wu H, Shinkai Y, Irizarry RA, and Feinberg AP. 2009. Large organized chromatin K9-modifications (LOCKs) distinguish differentiated from embryonic stem cells. Nature Genetics 41: 246.

Whyte WA, Orlando DA, Hnisz D, Abraham BJ, Lin CY, Kagey MH, Rahl PB, Lee TI, and Young RA. 2013. Master transcription factors and mediator establish super-enhancers at key cell identity genes. Cell 153: 307–319.

Zhang Y, Liu T, Meyer CA, Eeckhoute J, Johnson DS, Bernstein BE, Nusbaum C, Myers RM, Brown M, Li W, et al.. 2008. Model-based analysis of ChIP-Seq (MACS). Genome Biology 9: R137.

